# Generation of Arctic-like Rabies Viruses Containing Chimeric Glycoproteins Enables Serological Potency Studies

**DOI:** 10.1101/150300

**Authors:** Emma M. Bentley, Ruqiyo Ali, Daniel L. Horton, Davide Corti, Ashley C. Banyard, Anthony R. Fooks, Edward Wright

## Abstract

Rabies viruses have the highest case fatality rate of any known virus and are responsible for an estimated 60,000 deaths each year. This is despite the fact that there are highly efficacious vaccines and postexposure prophylaxis available. However, while it is assumed these biologics provide protection against all rabies virus isolates, there are certain subdivisions of RABV lineages, such as within the Arctic-like RABV (AL rabies virus lineage, where data is limited and thus the potency of existing biologics has not been thoroughly assessed. By fusing the Arctic-like rabies virus envelope glycoprotein ecto- and transmembrane domains with the vesicular stomatitis virus cytoplasmic domain, a high titre (7.7 x 10^5^ − 6.1 x 10^6^ RLU/ml) pseudotyped virus was generated that was subsequently used in a pseudotyped virus neutralisation assay. These results showed that Arctic-like rabies viruses are neutralised to human, canine and feline vaccines and human post-exposure prophylaxis and this was not influenced by the swapping of the cytoplasmic domains (CVS-11 vs CVS-11etmVSVc; *r* = 0.99, *p* < 0.0001). This study supports the concept that rabies virus vaccines and newly identified mAbs are able to neutralise rabies virus variants that cluster in a monophyletic clade, referred to as phylogroup I lyssaviruses.

## INTRODUCTION

Rabies, a neglected zoonotic disease caused by members of the *Lyssavirus* genus, poses a significant public health threat with a near 100% case fatality rate in infected individuals who do not receive pre or post-exposure prophylaxes (Fooks *et al.*, 2014). Globally, rabies virus (RABV) is accountable for an estimated 60,000 human deaths per year and the highest mortality rate of any other zoonotic disease when licenced vaccines and post-exposure prophylaxes are not administered (Fooks *et al.*, 2014). Serological studies, monitoring responses to pre- and post-exposure prophylaxis and undertaking widespread sero-surveillance, are vital aspects for the implementation of control programmes to lower rabies incidence (Banyard *et al.*, 2013; Brookes *et al.*, 2005; Wright *et al.*, 2009). However as many rabies-endemic areas are in the developing world, countries lack the infrastructure to be able to undertake these routine serological techniques (Banyard *et al.,* 2013).

Fourteen members of the *Lyssavirus* species are classified, with RABV being the prototype species (Dietzgen *et al.*, 2011). Two putative members of the *Lyssavirus* genus remain to be characterised and officially classified, Lleida bat lyssavirus (Ceballos *et al.*, 2013) and Gannoruwa bat lyssavirus (Gunawardena *et al.*, 2016). As infection with each species of the *Lyssavirus* genus causes a clinically indistinguishable disease the true burden of death from *Lyssavirus* species other than classical RABV is undefined. Arctic-like rabies viruses (AL RABV) form one of up to eight potential geographically and genetically distinct viral lineages of the RABV species (Kuzmin *et al.,* 2008; Mansfield *et al.*, 2006; Nadin-Davis *et al.*, 2007; Troupin *et al.*, 2016). Endemic across the Middle East and Asia, AL RABV is likely responsible for a significant proportion of rabies cases in these regions, which is proposed to result in greater than 20,000 human fatalities each year in India alone (Sudarshan *et al.*, 2007). Yet due to inadequate reporting systems and a weak healthcare infrastructure across this region, it is likely the true burden of rabies is far higher (Banyard *et al.*, 2013; Pant *et al.*, 2013). This lack of accurate data has led to the low prioritisation of control programmes by policy makers and public health professionals (Fooks *et al.*, 2014; Sudarshan *et al.*, 2007). Domestic dogs are the predominant transmission vector in human cases, and as the annual economic cost of canine rabies alone is estimated to be 8.6 billion USD, the economic and societal implications of endemic rabies are severe (Hampson *et al.*, 2015; Pant *et al.*, 2013). Along with the implementation of control programmes to limit rabies incidence, it is also important to undertake a comprehensive analysis of currently circulating RABVs and monitor for the emergence of new variants (Matsumoto *et al.*, 2013). While there is no evidence to indicate that AL RABV has an altered pathogenicity the protection afforded by vaccines and antivirals has not been specifically addressed. Thus it is important to fully understand the public health threat posed by this AL RABV lineage.

Pseudotyped virus (PV), a replication defective viral particle acting as a surrogate for live virus, has been used in a range of applications including serological assays and as vaccine immunogens (Mather *et al.*, 2013; Temperton *et al.*, 2015). The development of a PV neutralisation assay (PVNA) for the measurement of anti-rabies virus neutralising antibodies (VnAbs) in vaccine recipients, along with further large scale in-field sero-surveillance within a developing country has previously been described, providing sensitive and specific results which correlate with live virus assays and distinguishes between lyssavirus species (Temperton *et al.,* 2015; Wright *et al.,* 2008, 2009, 2010). As the use of PV allows neutralisation assays to be undertaken in biosafety level 1 or 2 laboratories, along with having a lower cost implication, the serological study of rabies can be expanded to resource-limited laboratories in regions where the virus is endemic.

While *Lyssavirus* isolates have previously pseudotyped efficiently, AL RABV pseudotypes fail to generate an adequate titre to allow use in downstream neutralisation assay studies. The flexibility of using a chimeric glycoprotein to produce recombinant, live-virus RABV has previously been demonstrated (Foley *et al.,* 2000). This was further applied in a study showing pseudotyping efficiency could be increased by altering the envelope glycoprotein, replacing the cytoplasmic domain with that of a glycoprotein that pseudotypes highly effectively(Carpentier *et al.,* 2011). This study has adapted this approach to produce a chimeric AL RABV envelope glycoprotein, increasing PV titre, and undertaking PVNA to assess vaccine and antiviral efficacy against this AL RABV lineage.

## RESULTS

### Chimeric AL RABV envelope glycoprotein construction and PV titre

Chimeric envelope G constructs were generated for four AL RABV isolates (RV61, RV193, RV250 and RV277), selected based on clinical significance (RV61) and reported poor growth in reference laboratory live viral cultures (RV193 and RV277; APHA, UK) and to represent three genetically distinct clades of the Arctic-related lineage (Fig. 1). The cytoplasmic domain sequence of each G was replaced with that of CVS-11 or VSV G. The ecto-transmembrane (etm) domain was not altered. A chimeric CVS-11 G sequence with a VSV G cytoplasmic domain (CVS-11etmVSVc) was produced for use as an internal control. Fig. 2 depicts the chimeric envelope G sequences generated.

**Fig. 1.**
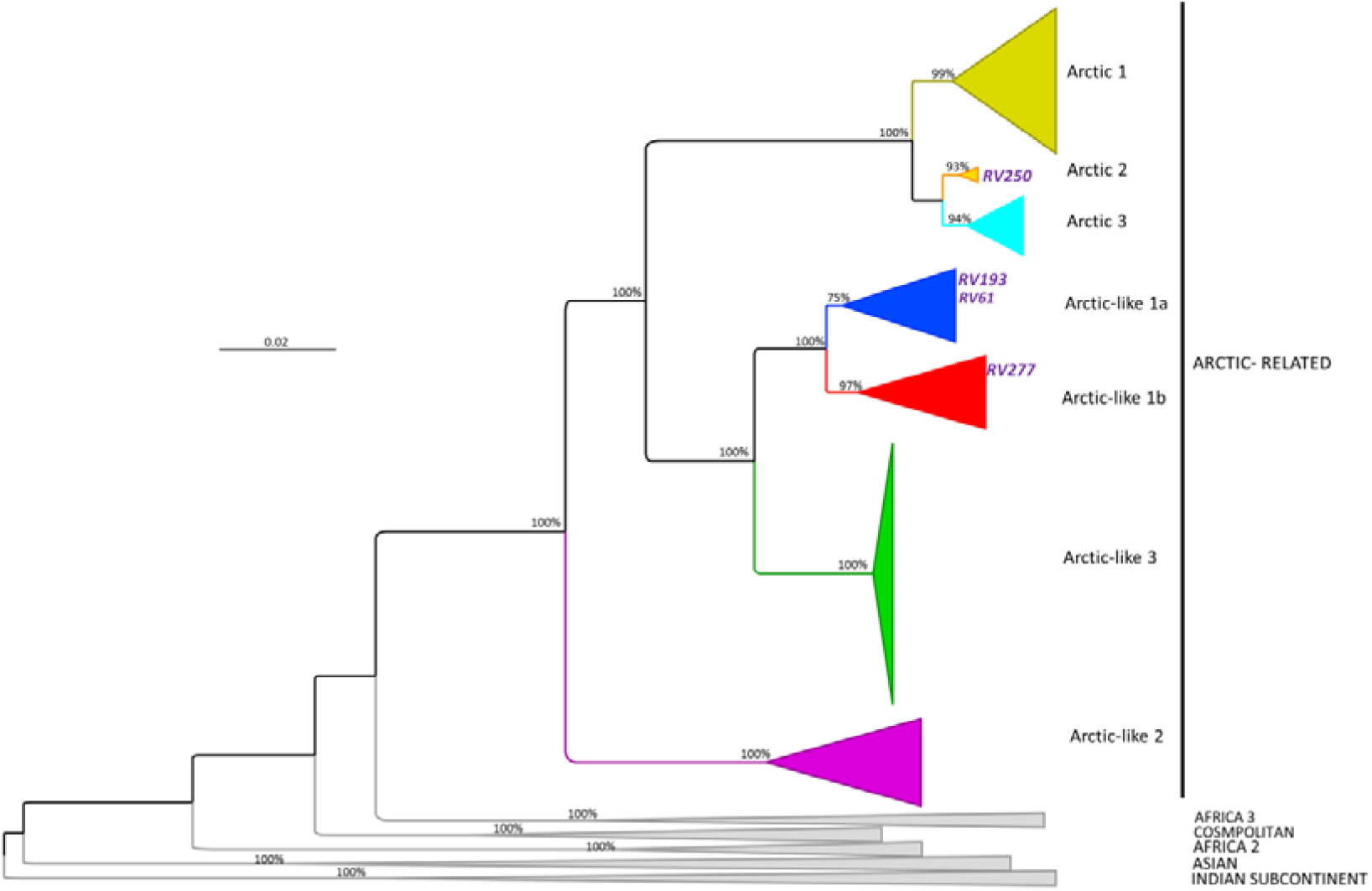
Maximum likelihood phylogenetic tree of 96 RABV glycoprotein coding sequences. Branches are labelled with bootstrap values at key nodes. Established clade, sub-clade and lineages are illustrated as previously defined (Pant *et al.,* 2013) and all except the Arctic-related viruses are collapsed for clarity. Positions of the viruses used in this study (RV61, RV193, RV250 and RV277) are indicated.

**Fig. 2.**
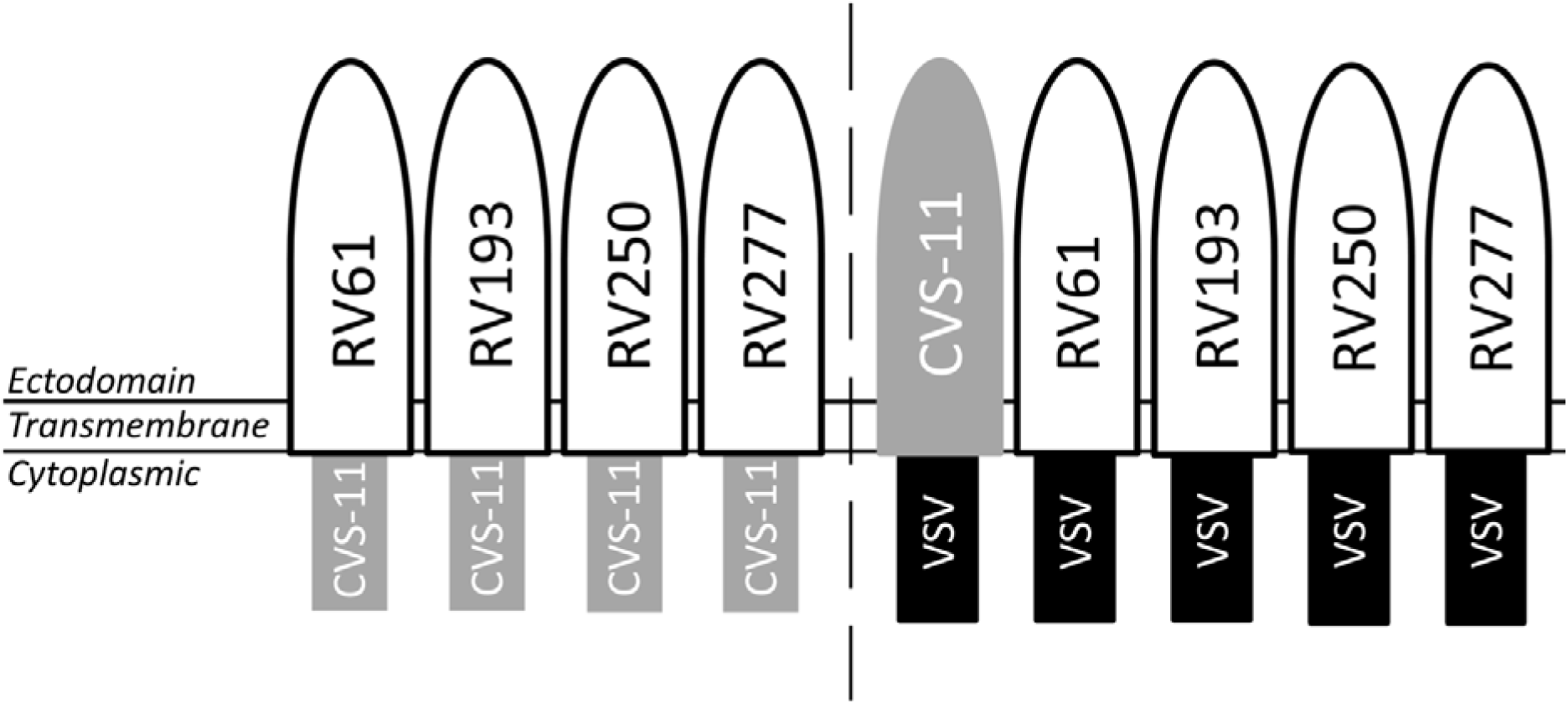
Schematic of the chimeric envelope glycoprotein constructs generated. AL RABV and CVS-11 ecto-transmembrane domains span from amino acid 1-480 of the full length G and the CVS-11 cytoplasmic domain from amino acid 481-526. The VSV G cytoplasmic domain used was amino acids 483-512.

Lentiviral PV comprising the wildtype and chimeric envelope G constructs were produced, packaging either an emerald green fluorescent protein (emGFP) or firefly luciferase reporter gene by transfecting HEK 293T/17 cells. These were titrated onto permissive BHK-21 cells to determine the viral titres of the chimeric constructs in comparison to that achieved using wildtype envelope G. Luciferase reporter PV with a chimeric CVS-11 cytoplasmic domain envelope G caused a decrease (−15.7 fold, *p* = 0.2; −10.6 fold, *p* = 0.0007; −1.4 fold, *p =* 0.7) or insignificant increase (1.2 fold; *p =* 0.7) in viral titre in comparison to the wildtype envelope G (Fig. 3). However, PV with a chimeric VSV cytoplasmic domain envelope G resulted in a significant increase (11.3 to 83.3 fold; *p* < 0.0005) in viral titre for three of the AL RABV isolates and a small (1.1 fold; *p* = 0.3) increase in viral titre for the RV250 isolate (Fig. 3).

**Fig. 3.**
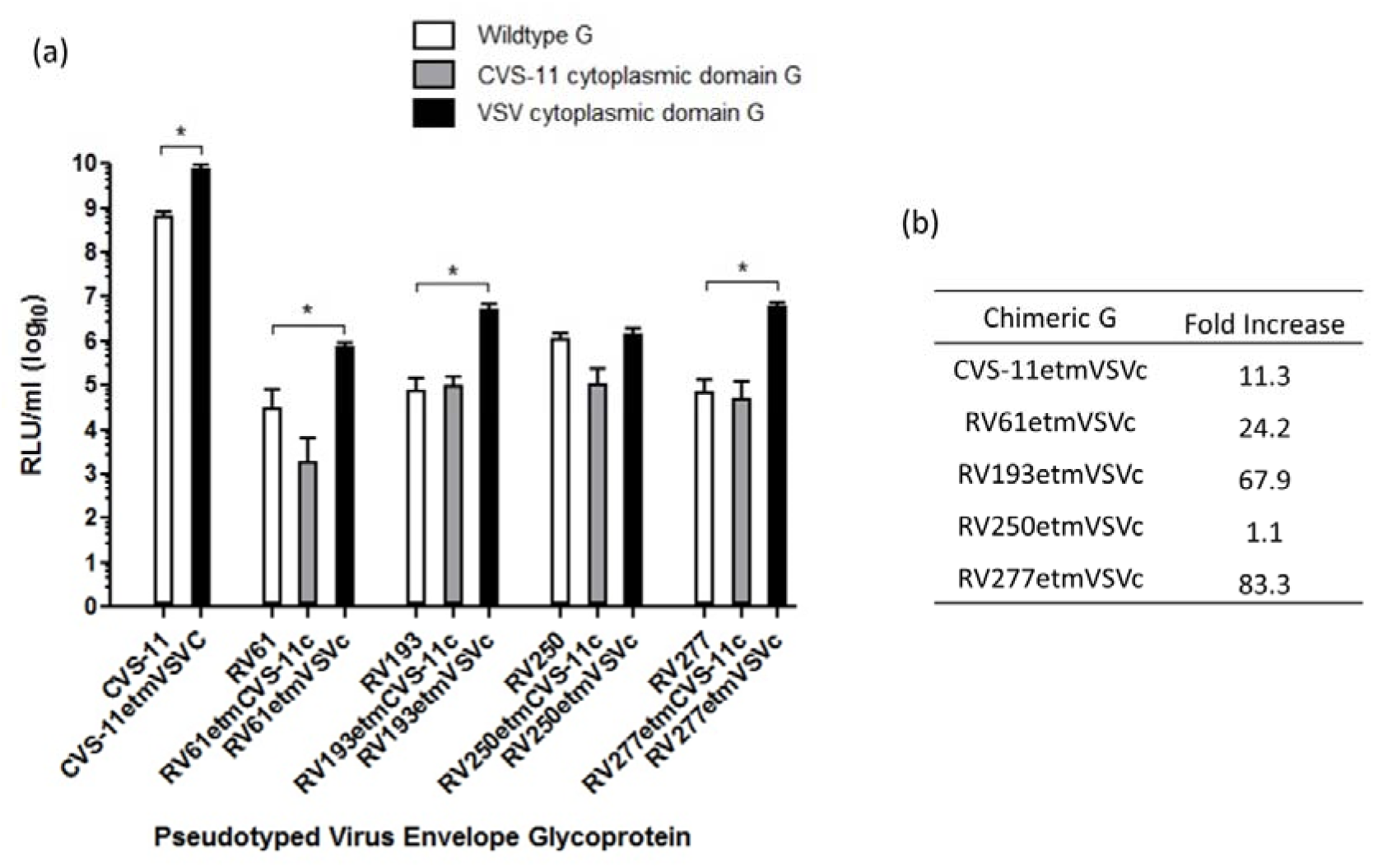
Comparison of viral titres using wildtype and chimeric envelope glycoproteins, (a) Aliquots of PV with a luciferase reporter gene were titrated on BHK cells to determine if a chimeric envelope glycoprotein with a CVS-11 or VSV cytoplasmic domain (CVS-11c or VSVc) increased titre, measured in relative light units per ml (RLU/ml) (\**p* < 0.0005; two-tailed *t*-test). Error bars show SD. (b) Fold increase in viral titre calculated from RLU/ml in (a) for chimeric VSVc G PV compared to wildtype G PV.

To corroborate this observation, PV with an emGFP reporter gene were generated and used in similar infection studies. Importantly, the increase in titre was also observed by fluorescent microscopy (Fig. 4a) and the fold increase determined by flow cytometry (Fig. 4b) was in line with that observed for the luciferase reporter PV constructs.

**Fig. 4.**
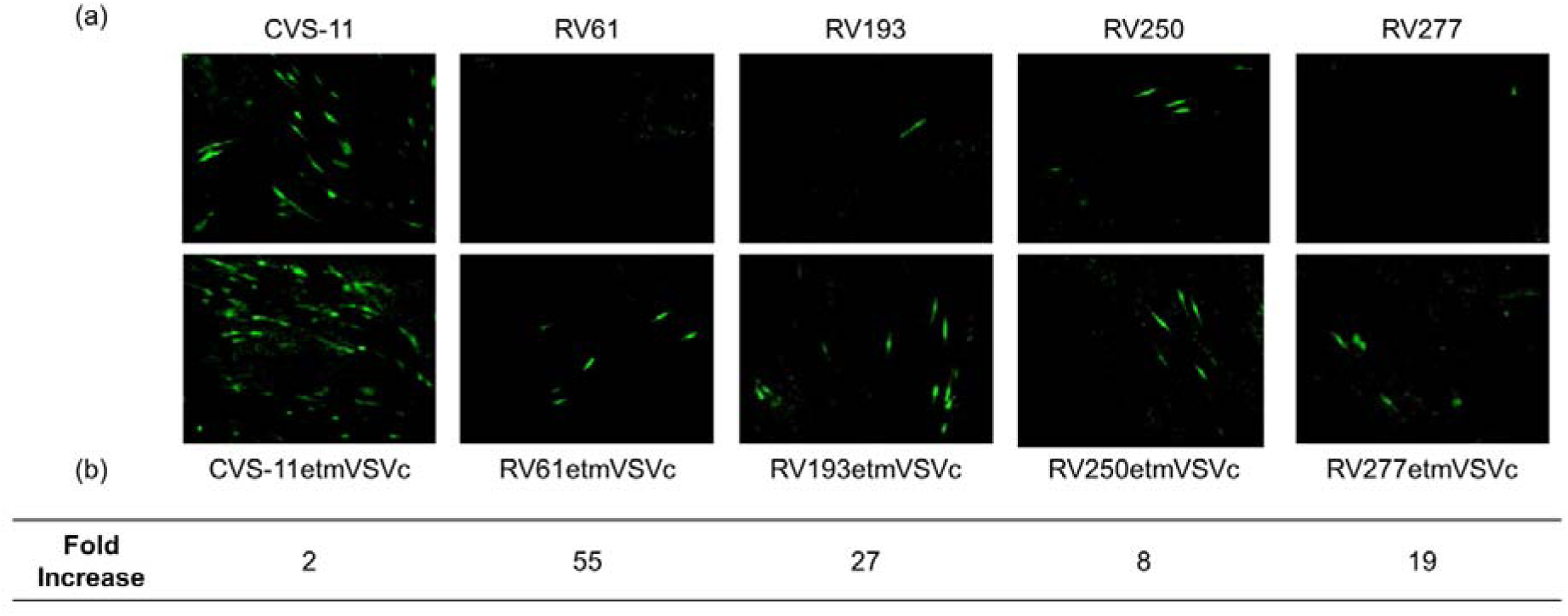
Viral titre comparison of PV bearing chimeric envelope glycoprotein and carrying an emGFP reporter gene, (a) Fluorescent micrographs of BHK cells infected with wildtype or chimeric VSVc G PV. (b) Fold increase in viral titre comparing chimeric VSVc G PV to wildtype stocks used in (a), determined by flow cytometry analysis.

### Current vaccines and biologicals neutralise Arctic-like rabies virus

The increased PV titre achieved for the AL RABV isolates using a chimeric VSV cytoplasmic domain envelope G enabled serology studies to be undertaken via a PVNA, to assess the level of sero-conversion afforded by current vaccines and post-exposure prophylaxis. The receptor-binding domain and antigenic sites of the RABV envelope G have been mapped to the ectodomain (Evans *et al.*, 2012; Kuzmina *et al.*, 2013); consequently switching the cytoplasmic domain to generate chimeric constructs should not influence the serological profile. Sequence comparison of etm domains of the AL RABV isolates and CVS-11 G shows high homology (Fig. 5). This analysis suggests the neutralisation profiles should be similar, yet as it is based on sequences alone it only serves as a crude estimate due to the disproportionate effects of individual amino acids on antigenic properties.

**Fig. 5.**
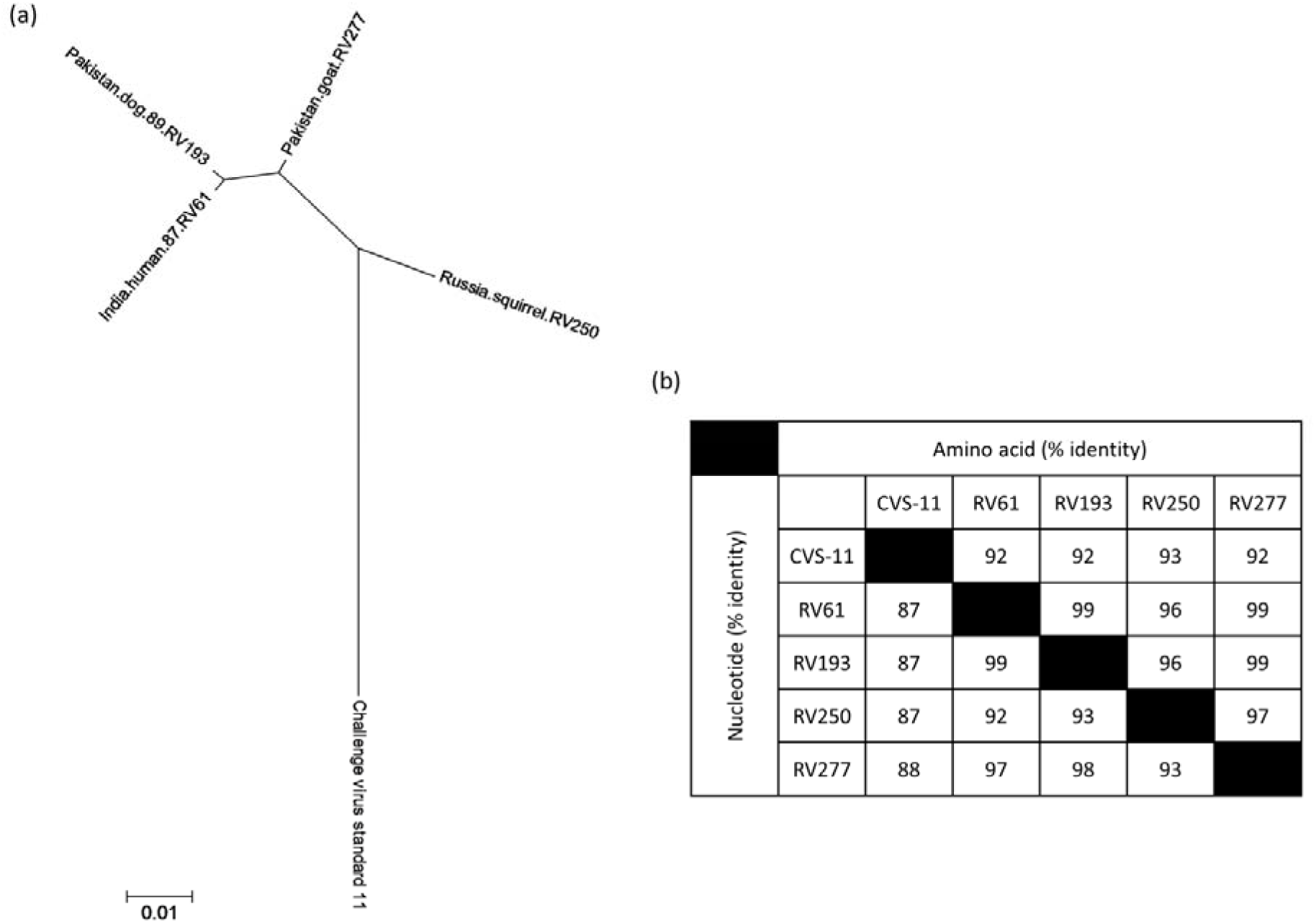
Degree of nucleotide and amino acid sequence identity between envelope glycoprotein etm domains of the rabies virus isolates within this study. (a) The radial phylogenetic tree scale corresponds to amino acid substitutions per site, (b) Nucleotide and amino acid percentage identities are shown.

Initially, the chimeric envelope G PV were tested alongside wildtype CVS-11 G PV using the OIE standard reference dog serum at a concentration of 0.5 IU/ml and WHO 2^nd^ international human anti-rabies Ig reference serum (2 IU/ml; prepared by NIBSC, UK) over a total of twelve doubling dilutions. Chimeric CVS-11etmVSVc G PV recorded an IC_100_ titre matching (IC_100_ = 80) or within one doubling dilution (IC_100_ = 269) of that for wildtype CVS-11 G PV for the OIE and WHO standards respectively (Fig. 6). Further to this, all AL RABV chimeric envelope G PV were neutralised at an equivalent or more potent level by each standard than that recorded for CVS-11 G PV.

**Fig. 6.**
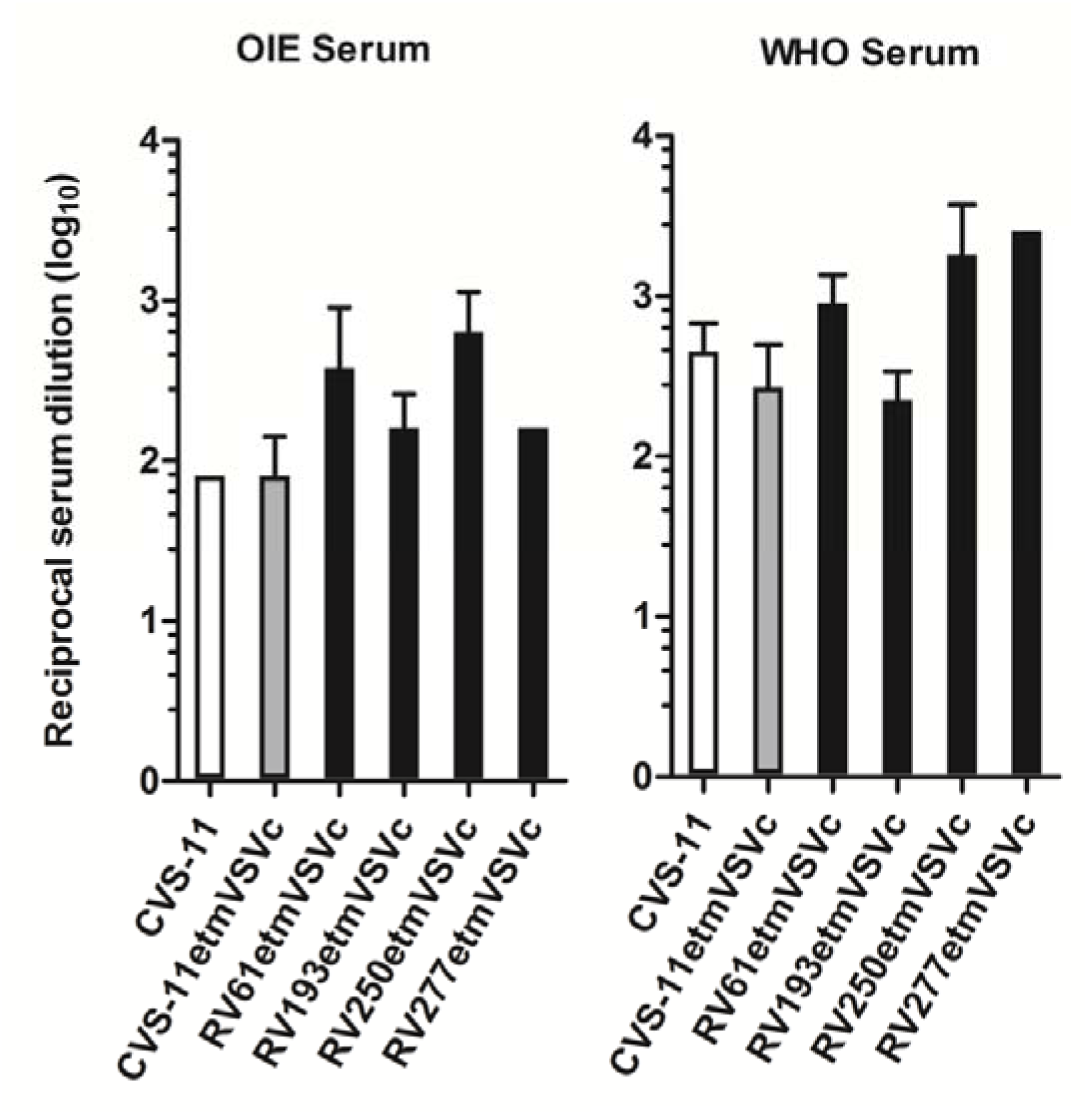
Neutralisation of PV by OIE and WHO serum standards. The OIE is a standard reference dog serum at 0.5 IU/ml and the WHO is the 2^nd^ international human anti-rabies Ig reference serum at 2 IU/ml. Values are reported as IC_100_ endpoint reciprocal dilutions (geometric mean ± SD). Where error bars are absent, replicates produced the same IC_100_ endpoint dilution.

Analysis of the neutralisation afforded against these AL RABV isolates by pre-exposure vaccination was undertaken by assessing a blinded panel of serum samples taken from RABV-vaccinated humans and domestic animals (dogs and cats) vaccinated as part of the UK PETS. The samples had previously been given a titre (IU/ml) using the fluorescent antibody virus neutralisation (FAVN) test method for detecting rabies specific antibodies, a score of 0.5 IU/ml by FAVN is considered the cut-off for adequate sero-conversion for protection (Cliquet *et al.*, 1998; WHO, 2013). When un-blinded, four human serum samples (H1, H5, H6, H7) with VnAb levels of 0.03 – 0.1 IU/ml, did not neutralise any PV tested (data not shown) and one sample with a VnAb level just below 0.5 IU/ml (H61, 0.38 IU/ml) neutralised each of the PVs (Fig. 7a). All samples with a VnAb titre above 0.5 IU/ml produced high levels of neutralisation for the CVS-11 and CVS-11etmVSVc G PV (IC_100_titres of 160-640) along with comparable levels for the AL RABV PV (Fig. 7a). The same cut-off is used to assign a satisfactory vaccination response in canine and feline recipients. All adequate animal serum samples produced a strong neutralising response (Fig. 7b). Of the four samples tested which had previously demonstrated VnAbs titres between 0.07 – 0.38 IU/ml on FAVN testing (Fig. 7b; PET-5531,-5545,-5734,-5896) a low level of PV neutralisation was detected (IC_100_titres of 12-57).

**Fig. 7.**
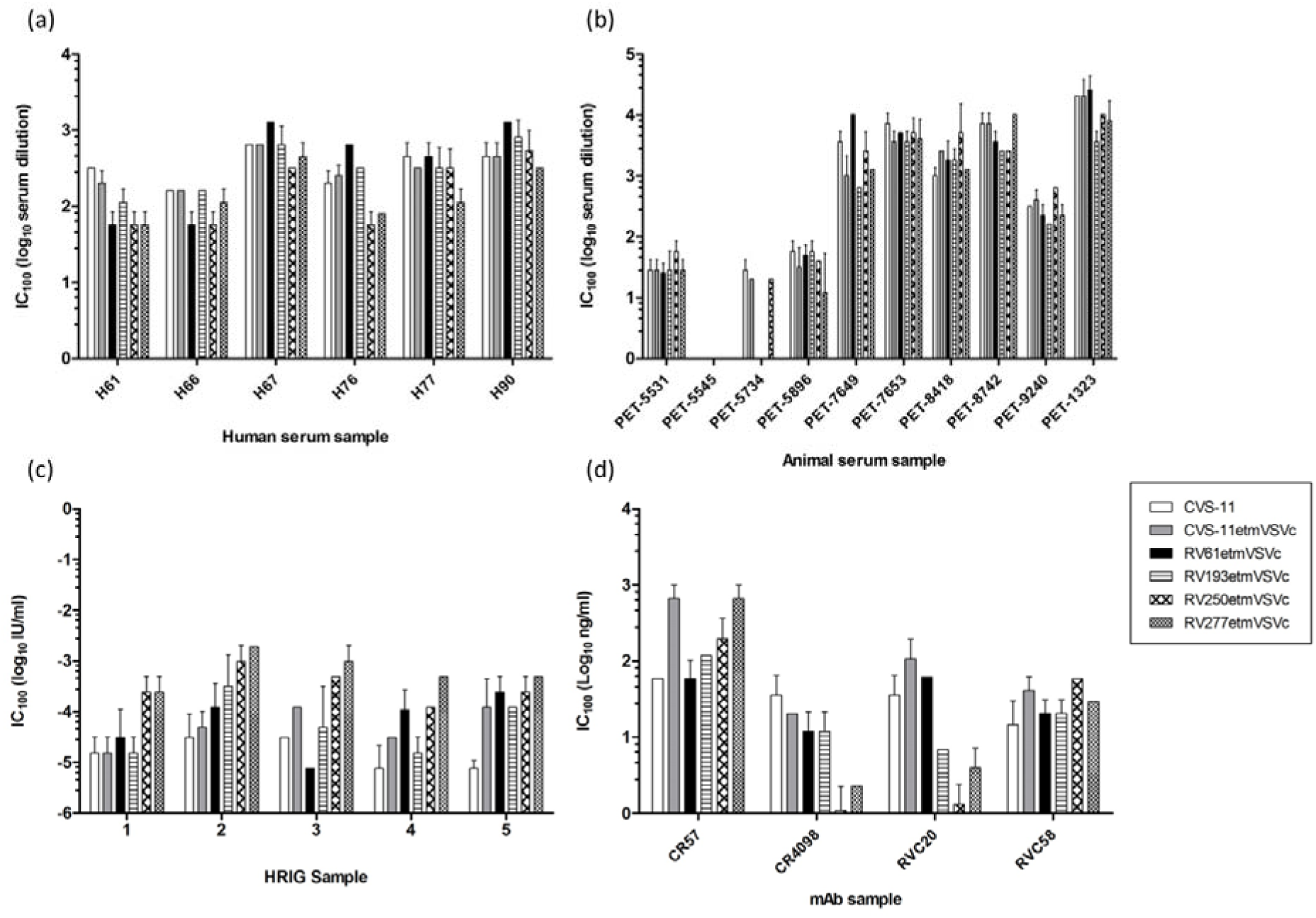
Neutralisation IC_100_ endpoint dilutions for human and animal serum samples, HRIG and mAb samples are reported for CVS-11 and AL RABV G PV. (a) Human serum samples are from RABV vaccine recipients, sample H61 has a titre of 0.38 IU/ml and the remaining samples a titre > 0.5 IU/ml. IC_100_ values are reported as reciprocal serum dilutions, (b) Animal serum samples are from vaccinated dogs or cats, four samples with titres between 0.07 − 0.38 IU/ml (PET-5531,-5545,-5734,-5896} are shown. The remaining samples have a titre > 0.5 IU/ml and IC_100_ values are reported as reciprocal serum dilutions, (c) HRIG samples were tested with a starting concentration of 2 IU/ml. (d) mAb samples are derived against different neutralising epitopes and were used at a starting concentration of 15 μg/ml. All values are reported as the geometric mean ± SD and where error bars are absent, replicates produced the same IC_100_ endpoint dilution.

Biologics used for post-exposure prophylaxis (PEP) were shown to effectively neutralise all AL RABV PV. Human rabies immunoglobulin (HRIG) samples were tested with a starting concentration of 2 IU/ml, each sample provided a potent level of neutralisation (Fig. 7c). Monoclonal antibody preparations, directed against various neutralising antigenic sites on the RABV envelope G and being considered for development to replace HRIG in PEP (Bakker *et al.,* 2005), were used at a starting concentration of 15 μg/ml. Each mAb neutralised the AL RABV isolates (IC_100_titre of 1.1 – 662.3 ng/ml), with CR4098 and RVC20 offering the most potent levels of neutralisation across all PV preparations (IC_100_titres between 1.1 – 35.6 and 1.3 – 106.9 ng/ml respectively; Fig. 7d).

The influence of switching the cytoplasmic domain on the neutralisation profile was also assessed by correlating IC_100_ titres obtained by PVNA for wildtype CVS-11 G PV alongside those for chimeric CVS-11etmVSVc G PV (Fig. 8). A strong correlation was shown between the PVNA results (*r* = 0.99, *p* <0.0001 [Pearson’s correlation]) thus switching the cytoplasmic domain does not alter the antigenicity of the envelope G.

**Fig. 8.**
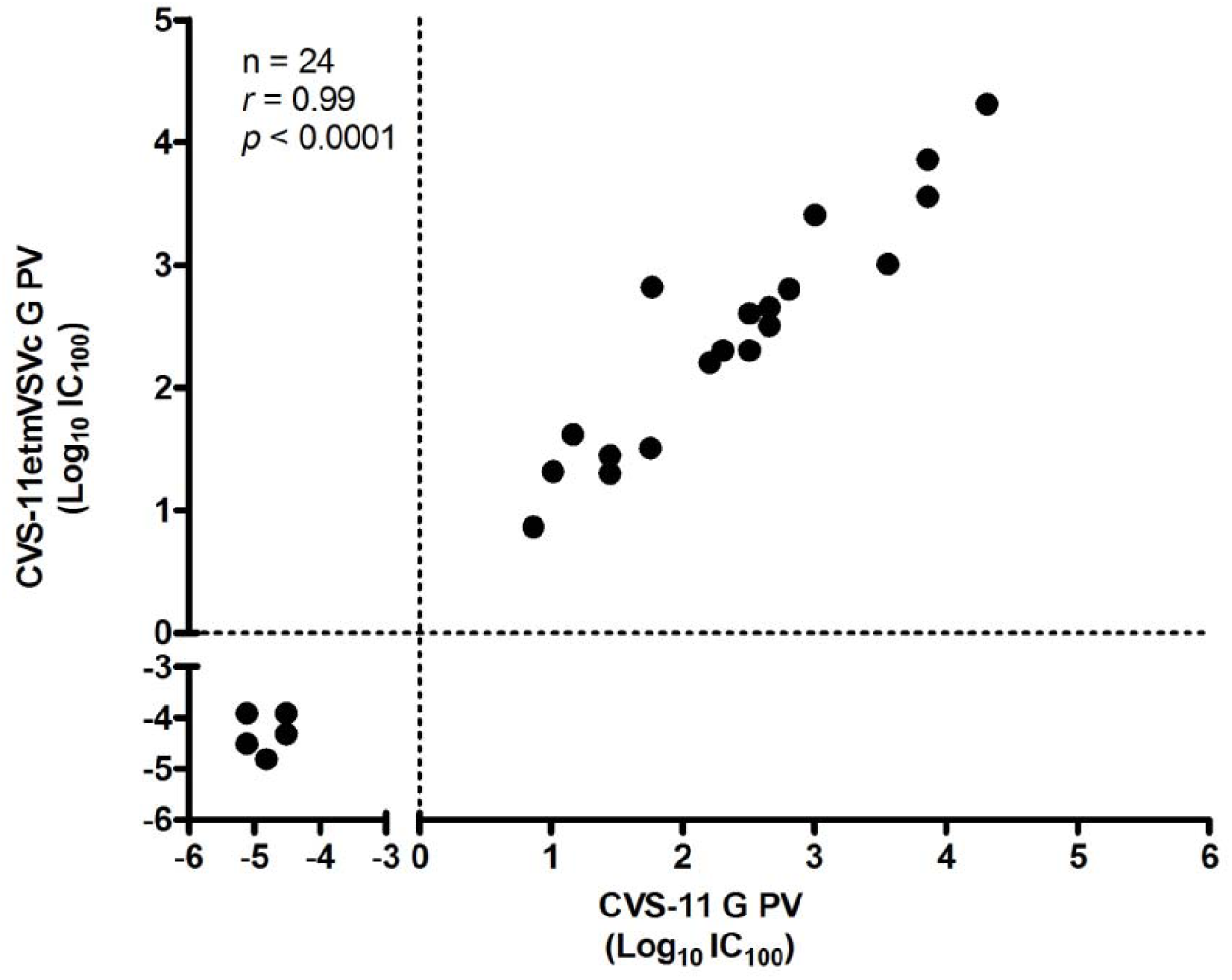
Comparison of the neutralisation IC_100_ endpoint for wildtype CVS-11 G PV compared to chimeric CVS-11etmVSVc G PV. A high correlation (*r*) was observed. Pearson’s product-moment correlation was used to calculate *r* and *p* values.

## DISCUSSION

Serological studies are required to define VnAb titres as part of vaccination and antiviral development and treatment schedules, while also allowing surveillance of the epidemiological spread of emerging viruses. As PV incorporate envelope proteins identical to the wildtype virus in their envelope they are antigenically similar, mimic the action of live virus in neutralisation tests and have proven to be a safe, robust and flexible alternative for use in serological assays (Mather *et al.*, 2013; Temperton *et al.*, 2015). The PVNA can be undertaken in containment level 1 and 2 laboratories as it does not require the handling of live virus, as opposed to other rabies virus neutralisation tests. The range of reporter genes and removal of the need for cold-chain storage make the PVNA an accessible and lower cost alternative to conventional techniques. Further to this, using a CVS-11 G PV, the PVNA proved to be 100% specific and equally sensitive to the WHO and OIE endorsed FAVN method of rabies VnAb detection (Wright *et al.,* 2008, 2009). This study further substantiates its use by demonstrating the inherent flexibility of the platform, allowing manipulation of the envelope G to increase PV titre, permitting serological studies to determine the protection conferred by vaccines and antivirals against AL RABV isolates. While the PVNA is primarily a research tool at this time, it has previously been used in clinical trials to assess vaccines (Ewer *et al.,* 2016; Ledgerwood *et al.,* 2010).

Chimeric AL RABV envelope G sequences were constructed with either a CVS-11 or VSV G cytoplasmic domain in an attempt to increase PV titre. Both CVS-11 and VSV G routinely produce high titre PV. Only the chimeric VSV cytoplasmic domain envelope G resulted in a significant increase in PV titre for three of the AL RABV isolates. The lower increase in titre for the RV250 isolate is thought to be attributed to a difference in its glycoprotein structure, phylogenetic analysis showed greater sequence homology between the other isolates which formed a separate cluster. Previously, the use of a chimeric CVS (B2c strain) envelope G with a VSV cytoplasmic domain was described (Carpentier *et al.,* 2011), reporting a two fold increase in titre; matching that observed for the CVS-11etmVSVc G used within this study. The mechanism behind this effect remains to be fully elucidated, yet several studies have described that the assembly of viable virions requires a direct or indirect interaction between the lentiviral matrix protein and envelope protein cytoplasmic domain (Cosson, 1996; Freed, 1998; Sandrin *et al.,* 2004; Yu *et al.,* 1992). Thus it is possible the cytoplasmic domain of VSV G interacts more effectively with the lentiviral matrix protein compared to that of CVS-11 G. Alternatively, it has been suggested a truncated or shorter cytoplasmic domain, as with VSV G, may cause a reduced steric hindrance or allow incorporation into lentiviral particles independent of matrix protein interaction (Freed & Martin, 1995). This is further supported by the report that truncation of the measles virus fusion (F) protein cytoplasmic domain lead to an increased PV titre (Frecha *et al.*, 2008).

Importantly, the regions of the VSV G defined within the literature differ due to the predictive nature of structural models. In this study, the VSV G cytoplasmic domain followed that used by Carpentier *et al.* (2011), as defined by Roche *et al.* (2006). However, caution is needed when designing chimeric sequences as alterations to some regions may result in loss of function, in particular the transmembrane domain which is involved in viral fusion and has more variability in the reported sequence (Cleverley & Lenard, 1998).

While current vaccines provide protection against RABV, the high level of sequence identity between the AL RABV isolates and CVS-11 G is not sufficient to definitively predict their neutralisation profile, as the effect of individual amino acid substitutions on antigenic variation has in some cases proven substantial (Horton *et al.*, 2010). Also, with the advent of mAbs for PEP, point mutations within the binding sites of mAbs can result in viral escape from neutralisation and thus the identification of these critical residues, assessing the neutralisation of generated escape viruses, forms a vital aspect in the development of effective, broadly neutralising, therapeutics (Bakker *et al.*, 2005; Marissen *et al.*, 2005). Direct measures of antigenic variation by serology are fundamental yet can prove difficult to quantify. The use of antigenic cartography has added power to the interpretation of antigenic data, enabling the generation of an antigenic map for a global panel of lyssaviruses, instrumental for predicting antigenicity based on the envelope G gene sequence (Horton *et al.*, 2010). The PVNA platform has previously been used in the collection of antigenic data in a cross-species comparison of lyssavirus neutralisation, showing suitability as a high-throughput screening method to complement quantification of antigenic differences (Wright *et al.*, 2008, 2009). This study further supports use of the PVNA, demonstrating inherent flexibility in the creation of chimeric viral envelope protein PV without disruption to the neutralisation profile and therefore the envelope protein function. This enabled confirmation of sero-conversion, and by extrapolation, protection afforded by current vaccines and prophylaxis against the AL RABV isolates.

The AL RABV isolates were found to be effectively neutralised by human and mammalian serum samples, conferring adequate protection by current pre-exposure vaccine formulations. As more than 99% of human rabies cases occur following contact with rabid dogs, the control of rabies within this population is of high priority (Banyard *et al.*, 2013; WHO, 2013). Mass vaccination campaigns targeting dog populations are highly effective and thus monitoring levels of protection afforded by animal vaccine formulations is of equal importance to the prevention of human rabies infections. All licenced vaccine preparations are derived from inactivated preparations of classical RABV, which has shown to confer protection against viruses in phylogroup I but offer limited or no protection against those in phylogroups II and III (Evans *et al.*, 2012; Fooks, 2004; Hanlon *et al.*, 2005). Since AL RABV is a lineage of classical RABV, the protection observed follows this accepted consensus and therefore, even though the isolates of rabies causing cases in the regions where AL RABV circulate are not fully characterised, unexplained vaccine failures have not been reported. However, due to poor growth of these AL RABV isolates in live viral cultures, which could suggest a different structure of the G protein, and the implication of one isolate in a transplant-associated rabies outbreak in Germany (Ross *et al.*, 2015), it was important to be able to undertake serological evaluation. Further studies into cross-protection of rabies vaccines against more divergent lyssaviruses, such as those within phylogroups II and III, using this PVNA could assist in the development of a more broadly cross reactive vaccine formulation.

PEP regimes have long been effective in preventing rabies virus infection in the event of exposure. For previously un-vaccinated individuals this consists of wound cleansing, vaccination and the administration of rabies immunoglobulin (RIG) to provide passive immunity in the interval before vaccine induced active immunity is achieved (Fooks *et al.*, 2014). RIG of human (H) or equine (E) origin is available. While HRIG is preferred due to its longer half-life, it is expensive compared to the more immunogenic ERIG, which limits its use in the developing world; yet both are in short supply (WHO, 2013). The AL RABV isolates were neutralised by all HRIG preparations, however alternative means of PEP are now being sought by the development of mAb cocktails. Here we tested four mAbs, RVC20 and CR57, and RVC58 and CR4098, which target antigenic site I and III respectively of the RABV G (Bakker *et al.,* 2005; De Benedictis *et al.,* 2016; Marissen *et al.,* 2005). In order to meet WHO guidelines, which suggest RABV PEP should contain at least two antibodies to lower the probability of immune escape, CR57 and CR4098 have been combined into the CL184 mAb cocktail and undergone phase II clinical trials (Bakker *et al,* 2008; Nagarajan *et al,* 2014; WHO, 2013). In this study, each mAb effectively neutralised the AL RABV isolates, which can further serve as an indication that both antigenic sites are highly conserved across the AL RABV lineage.

Ultimately, the flexibility of using PV demonstrated within this study can be further extended. The generation of antigenic escape mutant envelope protein for incorporation into the PV platform will enable evaluation of mAb cocktails undergoing development. Likewise, switching of epitopes between lyssavirus envelope G can allow further cross neutralisation studies to be undertaken, an important aspect in vaccine design. The ability to switch domains of the lyssavirus envelope G has already been explored, highlighting the potential for use in antigenic studies (Jallet *et al,* 1999). This will enable the level of protection afforded against other divergent lyssaviruses in phylogroup II and III to be evaluated, of great interest from a public health perspective due to their unknown disease burden.

Using the approach of generating a chimeric envelope glycoprotein with a VSV cytoplasmic domain resulted in high titre PV without affecting their neutralisation profile. These data also provide further evidence of the flexibility pseudotyped virus-based assays provide when undertaking serological studies of highly pathogenic viruses. In conclusion, it was determined the AL RABV isolates are neutralised by available vaccines and post-exposure prophylaxis.

## METHODS

### Cell lines

Human embryonic kidney 293T clone-17 cells (HEK 293T/17; ATCC CRL-11268) were used for PV production and subsequent titration and PVNA were undertaken using baby hamster kidney-21 clone-13 cells (BHK-21; ATCC CCL-10). Both cell lines were cultured in Dulbecco’s Modified Eagle Medium (DMEM) supplemented with 10% foetal calf serum (FCS) and 1% penicillin/streptomycin with 5% CO_2_.

### Viruses and cloning of chimeric envelope glycoprotein

The four AL RABV envelope glycoprotein (G) genes used in this study were amplified from viral RNA of India.human.87.RV61 (KU534939), Pakistan.dog.89.RV193 (KU534940), Russia.squirrel.RV250 (KU534941) and Pakistan.goat.RV277 (KU534942) and cloned into the pl.18 expression vector (Cox *et al.,* 2002). The challenge virus standard-11 (CVS-11) (Wright *et al.*, 2008) and vesicular stomatitis virus (VSV; a gift from Didier Trono, Adgene plasmid # 12259) G genes have previously been described and were used to produce control pseudotyped virus in this study.

Chimeric envelope G were generated by overlap extension polymerase chain reaction (PCR) (Heckman & Pease, 2007). Specific primers, designed based on the envelope G gene sequences, were used to separately amplify DNA fragments encoding the ecto- and transmembrane domain (etm) and cytoplasmic domain (c) portions of the chimeric constructs using the proofreading enzyme AccuPrime *Pfx* SuperMix (Life Technologies, UK). Primers are listed in Table S1. A further PCR was used to bring the entire open reading frame together utilising the overlapping complementary regions initially introduced. Once amplified, unique restriction sites introduced by the primers at the 5’ and 3’ ends were used to clone the chimeric sequences into the pl.18 expression plasmid. Clones containing the correct insert were identified by restriction enzyme digest and confirmed by Sanger sequencing.

### Pseudotyped virus production and titration

Lentiviral pseudotyped virus production followed the transfection protocol previously described (Wright *et al.*, 2009). Briefly, the HIV *gag-pol* construct p8.91 and firefly luciferase reporter construct pCSFLW or emerald green fluorescent protein (emGFP) reporter construct pCSemGW (kindly provided by University College London (UCL), UK) (Cubitt *et al.*, 1998) were transfected concurrently with plasmid expressing the appropriate envelope G into HEK 293T/17 cells using Fugene-6 (Promega, UK) or polyethylenimine (PEI) (Sigma, UK) transfection reagent. Supernatant was harvested 48 and 72h post-transfection and filtered through a 0.45μm filter, storing long-term at −80°C.

Titration of pseudotyped virus aliquots carrying the firefly luciferase reporter gene was performed by transducing BHK-21 cells in a 96-well plate (2 x 10^4^ cells/well) with serially diluted pseudotype supernatant in quadruplicate. Following 48h incubation, cell luminescence was read using the Bright-Glo assay (Promega, UK) and GloMax-Multi+ microplate luminometer (Promega, UK) with titres expressed as relative luminescence units per ml (RLU/ml) or 50% tissue culture infective dose per ml (TCID_50_/ml). Likewise, titration of virus with emGFP reporter gene involved transducing BHK-21 cells in duplicate with doubling dilutions of pseudotype supernatant. emGFP positive cells were visualised using a fluorescent microscope and counted using a Dako CyAn ADP cytometer (Beckman Coulter, UK).

### Serum and mAb samples

The World Organisation for Animal Health (OIE) standard dog reference serum at a concentration of 0.5 international units per ml (IU/ml) and WHO 2^nd^ international human anti-rabies Ig reference serum (2 IU/ml; prepared by National Institute for Biological Standards and Control (NIBSC), UK) were used as positive controls. A range of sera (n=20) from RABV-vaccinated humans (Rabipur, Novartis) and dogs and cats (Rabvac, Fort Dodge; Nobivac, Intervet; Rabisin, Merial; Quantum, Schering Plough) enrolled in the UK pet travel scheme (PETS) were used (Ramnial *et al.*, 2010). Human monoclonal antibody (mAb) samples were produced as described in (De Benedictis *et al.,* 2016) and commercial rabies immunoglobulin (RIG) released for the European market were kindly provided by NIBSC, UK.

All samples were titrated in 2-fold serial dilutions. All experiments were undertaken at least in duplicate, where the titre varied by more than one doubling dilution it was repeated and the geometric mean recorded, as per standard serological practice (Bresson *et al.*, 2006).

### Neutralisation assays

The pseudotyped virus TCID_50_ value was calculated using the end point method (Condit, 2001). In a 96-well plate 50 x TCID_50_ pseudotyped virus was incubated with sera in duplicate for 1 hr at 37°C (5% CO_2_) before the addition of 1 x 10^4^ BHK-21 cells. After a further 48 hrs incubation cell media was removed and a 50:50 mix of Bright-Glo reagent (Promega, UK) and fresh media added. Luciferase activity was detected on a GloMax-Multi+ microplate luminometer (Promega, UK) and IC_100_ end-point titres recorded.

### Phylogenetic analysis

Analysis of 96 RABV glycoprotein sequences (1575 nucleotides) was inferred using MEGA6, with a GTR substitution model, gamma distribution of rate variation sites a proportion of invariant sites (GTR+G+I). Established lineages were illustrated and all except the Arctic-related viruses collapsed for clarity. Bootstrap values (100 replicates) were illustrated at key nodes.

## ACKNOWLEDGEMENTS

We thank Greg Towers (UCL) for providing the pCSemGW plasmid and Giada Mattiuzzo (NIBSC) for supplying the RIG samples used within this study. This work was supported by the Department for Environment Food and Rural Affairs (Defra); Scottish Government and Welsh Government (grant number SE0431).

## CONFLICTS OF INTEREST

Davide Corti is employed by Humabs Biomed, which is developing rabies monoclonal antibodies. Ruqiyo Ali undertook this work while an MSc student at the University of Westminster but has since taken up employment at AstraZeneca.

---

**Table S1.**
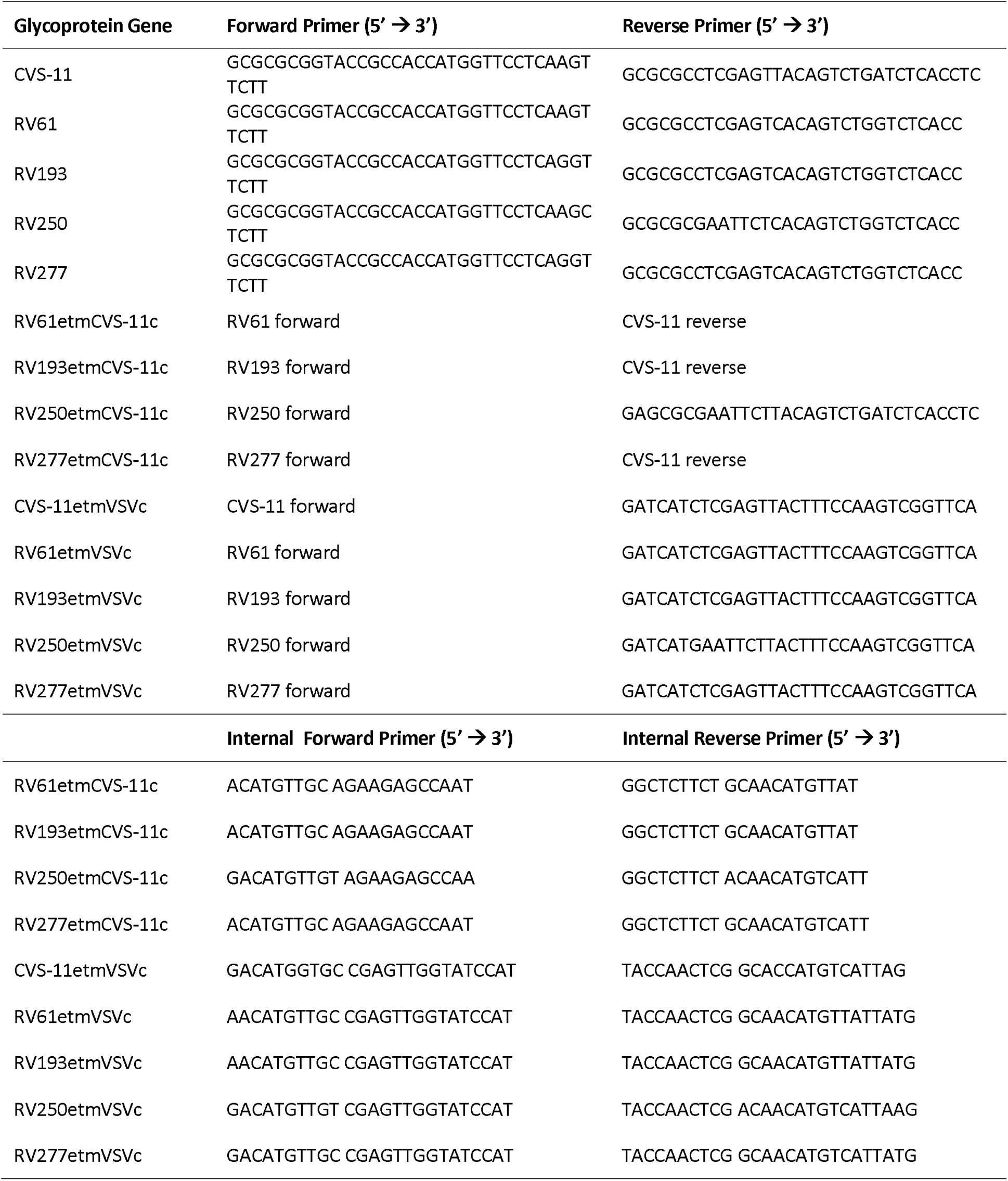
Oligonucleotide primers used for PCR amplification and overlap extension PCR to create chimeric envelope glycoprotein sequences.

